# Altered microRNA expression profles are involved in Storage Lesions of Apheresis Platelet

**DOI:** 10.1101/386011

**Authors:** Zhaohu Yuan, Qianyu Wu, Xiaojie Chen, Yaming Wei

## Abstract

Although platelet is anucleate cell, it contains a large amount miRNAs. This study aims to explore the relationship between miRNAs expression profles and platelets function, as well as miRNAs potential roles during platelet storage lesions. Platelets were collected from 15 healthy men with O blood types. MiRNAs profiles in platelets were detected by Agilent Human miRNA Array Differential miRNA levels were studied using human platelets after apheresis and stored for 2, 5 and 8 days using microarray. There were 167 and 230 altered miRNAs during the 5 and 8 day storage, respectively. In addition, the number of reduced miRNAs was much greater than that of increased. And many of them are involved in functions of platelet activation, degranulation, PDGF receptor signaling pathway and cell differentiation. The results of RT-PCR showed that the expression of miR-21-5p, miR-21-3p and miR-155 decreased on the 5th day, while miR-223, miR-3162 and let-7b increased. Flow cytometry results revealed that with increase of storage time, the expression of P2Y12 increased and phosphorylation level of VASP reduced. Meanwhile, the platelet reactivity index (PRI) decreased from 72.7% to 18.2%, while the apoptotic percentage of platelet significantly increased. For the first time, we found altered miRNAs are closely related to platelet aggregation, including P2Y12, VASP and GPⅡb/Ⅲa. Through KEGG database prediction, we verified there were many miRNAs impacting the pathway of platelet aggregation, such as miR-223, miR-21 and let-7b, which indicated miRNAs might serve as potential biomarkers of storage lesion in platelet. These target miRNAs are related to activation, degranulation and PDGF receptor signaling pathway of platelet.

## Introduction

Platelets are released into the blood circulation from the megakaryocytes of bone marrow as cytoplasmic fragments, and platelets are the second most abundant cells of the blood. The main role of platelets is to ensure primary hemostasis, which means the maintenance of blood vessel integrity and the rapid cessation of bleeding in the event of loss of vascular integrity [1, 2]. The formation of platelet is adjusted by thrombopoietin (TPO), platelet survival time in vitro is 8-11 days by 51 Chromium tag analysis [3]. According to the international standard of 1986, the duration of platelets in vitro preservation is five days [3, 4]. Due to the short validity of platelets, many researchers began to explore new ways to extend the shelf life of platelets. The newly developed PVC bag has extended the storage life of platelets to 7 days, and the storage life of frozen platelet treated with DMSO can be extended to one year [5]. With the technical progress of molecular biology, such as RT-PCR and RNA microarray analysis, the scientist began to explore function and mechanism of platelet in the molecular level to prolong the life-span in vitro.

MiRNAs are small noncoding RNAs endogenously involved in posttranscriptional regulation of cellular gene (messenger RNA, mRNA) expression [6, 7]. MiRNAs found in platelets are currently recognized as important regulators in the activation and aggregation of platelets [8]. Though the exact mechanism of miRANs is not yet clear, they combined to non-coding sequence of target gene 3′ UTR by the complementary base-pairing reactions to degrade or inhibit the translation of mRNA and participate in the regulation of mRNA transcription and post transcription [9]. With the development of sequencing technology, the study of miRNAs in platelet also progressed gradually. In the stored platelet, 52 miRNAs associated with apoptosis were analyzed, and yet only 4 of which were found with altered levels during the storage [10]. Another research suggested that the proportion of miR-127 and miR-320a served as a biological marker whether platelets could be continued to infuse or not [11]. But neither these studies are comprehensive nor showcase the complete miRNA changes of platelet during in vitro storage at 22 ± 2℃.

In this study, we analyzed the expression of 2007 miRNAs of apheresis platelets after 2, 5 and 8 days of storage by miRNA microarrays assay, some miRNAs and their predicted target genes were selected to validate by RT-PCR. MiRNAs expressions in apheresis platelet were altered during storage. Altered miRNAs are mainly related to platelet aggregation, miR-223, miR-21, let-7b and their target genes are related to platelet activation, degranulation, PDGF receptor signaling pathway and cell differentiation.

## Methods and Materials

### Ethics Statement

The study protocol received approval from the Ethics Committee of Guangzhou First People’s Hospital (K-2017-007-01). Written informed consent was obtained from all participants or their families.

### Participants

Apheresis platelets samples were collected from 15 healthy adult men with O blood type with median age 39.72 ± 2.43 years at Guangzhou First People’s Hospital, Guangzhou Medical University in 2017. The concentration of apheresis platelet is ≥0.9×10^9^/mL, WBC ≤1.0×10^4^/mL, RBC ≤2.00×10^7^/mL. Under normal procedure, platelet were apheresized and tested for qualified on the first day, the earlyest available day for clinic treatment is the 2nd, platelet is ready to use for patients in next 2-5 day, and 5 day is the longest day of storage ratified by CFDA, and 8 day is the theoretical longest platelet storage time in vitro. All procedures performed in studies involving participants were in accordance with the ethical standards of the Guangzhou First People’s Hospital.

### Total RNA Isolation

Total RNA was isolated using the Ribopure™ Blood RNA isolation Kit according to manufacturer’s protocol. Total RNA was isolated from apheresis platelets for further trials on 2nd, 5th and 8th day during storage. RNA concentration was determined using NanoDrop ND-2000 (Thermo Scientific, CA, USA). The integrity of RNA samples was verified using denaturing gel electrophoresis (15% polyacrylamide gel) and Agilent 2100 Bioanalyzer (Agilent Technologies, Santa Clara, CA, USA). Samples displaying RNA integrity number (RIN) of >7.5 were subsequently submitted for microarray and quantitative PCR analysis.

### RT-PCR assay

RT-PCR assay was performed as previously described (12). Briefly, 20 ng of total RNA was reverse transcribed (in 15 μL) using specific stem-loop primers. For the PCR reaction, 1.33 μL of RT-product was used. PCR was carried out using the Applied Biosystems 7900 high throughout sequence detection system (Applied Biosystems, Foster City, CA, USA). The RT-PCR-reactions were performed in triplicate, in three separate experiments. The small RNA U6, a control and normalizer for miRNAs, was relatively unchanged that excluded the possibility of artifactual changes in miRNA recovery. Gene expression analysis was performed as previously described citation [13] and the sequence of the primers are described in Table 1.

**Table 1.**
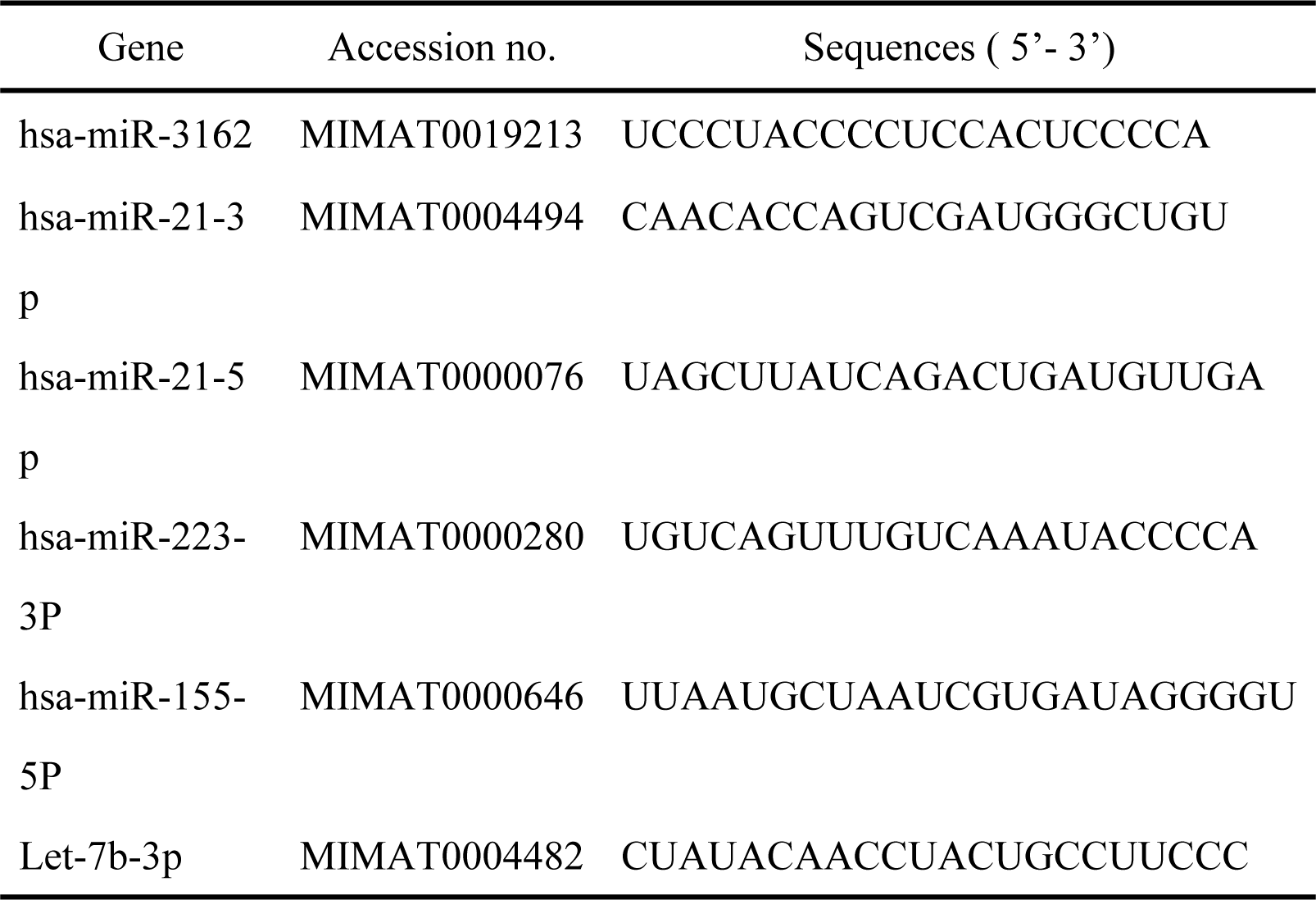
qPCR primer sequence

### RNA purification and microarray analysis of miRNA expression

Total RNAs were purified using Trizol-LS reagent (Thermo Scientific, Waltham, Massachusetts, USA) according to the manufacturer's protocol. The purity and integrity of the RNA sample was assessed using the ratio of absorbance at 230, 260 and 280 nm by NanoDrop ND-2000, a micro-volume spectrophotometer（ Thermo Scientific, USA） and Agilent 2100 Bioanalyzer ^®^ (Agilent Technologies, Santa Clara, CA, USA). Three random selected RNAs from above 15 samples were performed microarray analyses using SurePrint G3 Human V16 miRNA 8×60K microarray kit (Agilent Technologies), as previously described [14]. The labeled RNA was hybridized with the microarray kit in the Agilent Microarray Hybridization Chamber (Agilent Technologies, CA, USA) for 20 h. The fluorescence intensities of the labeled miRNA samples on the microarray were measured using Scanner and Feature extraction software (version10.7.1.1, Agilent Technologies, CA, USA). The digitalized fluorescence intensities were analyzed using GeneSpring GX version 12.5 (Agilent Technologies, CA, USA). List of miRNAs with an absolute fold change ≥2.0 and ≤0.5 and *p* value < 0.05 after the Benjamini-Hochberg false discovery rate (FDR) correction following multiple comparisons were considered significant [15].

### The prediction of target genes and bioinformatics analysis of miRNAs

To assess the biological functions of the altered miRNA expression, the bioinformatics analyses were performed to predict the target-related cellular pathways using the Kyoto encyclopedia of Genes and Genomes (KeGG) pathways and categorization of target genes with specific biological functions using the AmiGO Gene Ontology (GO) analysis tool (http://amigo.geneontology.org/amigo) [16]. The prediction of target genes was limited by setting the value of the threshold to 0.8 in the DIANA-microT tool. The KeGG pathway category was processed by setting the threshold of eASe score, a modified Fisher exact *p*-value, to 0.1, and involved KeGG pathways that displayed a value >1% (percentage of involved target genes/total target genes in each pathway) were selected. GO analysis was performed in eight categories for positive or negative regulation of apoptotic processes, cell growth, cell proliferation and cell cycle [17].

### Apoptosis assays by Flow cytometry

Phosphatidylserine (PS) exposure was measured using the binding of fluorescein-isothiocyanate-labeled (FITC) Annexin-V to PS according to the manufacturer's instruction. Briefly, platelets (2×10^6^ per mL) were washed in PBS and suspended in 500 µL binding buffer. Annexin V FITC (5 µL) and propidium iodide (5µL) were added to the cells for 10-15 minutes in the dark at room temperature and analyzed with FACSCalibur flow cytometer (Becton Dickinson, CA, USA) equipped with argon laser (488 nm) and filtered at 530 and 585 nm for FITC and phycoerythrin respectively. Low fluorescence debris and necrotic platelets, permeable to propidium iodide, were gated out before to analysis and 10^4^ events were collected [18]. Data were analyzed using Cell Quest software (Becton Dickinson, CA, USA). This assay was also used to measure spontaneous apoptosis of platelets.

### Analysis of VASP/P2Y12 phosphorylation by flow cytometry

To determine the VASP phosphorylation state of platelet, we used a standardized flow cytometric assay (Platelet VASP; Diagnostic Stago (Biocytex), France), which is an adaptation of the method of Schwarz et al. [19]. Briefly, platelets were permeabilized with non-ionic detergent, incubated with PGE1 or with PGE1 and ADP for 10 minutes and then fixed with paraformaldehyde. Platelets were labeled with a primary monoclonal antibody against serine 239-phosphorylated VASP (16C2), then followed by a secondary fluorescein isothiocyanate-conjugated polyclonal goat antimouse antibody. The total test duration did not exceed 30 min. Analysis were performed on a FACSCalibur flow cytometer (Becton Dickinson, CA, USA) at a medium rate, the population of platelet was identified from its forward and side scatter distribution. A platelet reactivity index (PRI) was calculated from the median fluorescence intensity (MFI) of samples incubated with PGE1 or PGE1 and ADP [20, 21].

### Statistics

All graphs were generated using Prism 4 (GraphPad Software, Inc., CA, USA). Statistical significance was assessed by using the oneway ANOVA test (for parametric data) or Kruskal–Wallis test (for nonparametric data) using SPSS 18.0 (SPSS, Inc., Chicago, IL, USA). When the *p* values were statistically significant, posthoc pairwise comparisons with the Tukey Honest Significant Difference (HSD) method were performed. Columns are the mean of triplicate experiments; bars ± SD. *p* < 0.05 was considered to be statistical significant.

## Results

### Differential miRNA expression levels during storage

By miRNA microarrays assay we analyzed the expression of 2007 miRNAs in apheresis platelets after 2, 5 and 8 days of storage. In this study, we compared the expression profles of miRNA in banked platelet between the 5th and 2nd day. As shown in Figure 1, 47 and 120 miRNAs are up- and down-regulated more than 2-fold, respectively. On the 8th day, total 230 miRNAs expression changed, 28 of them up-regulated, 202 of them down-regulated compared to 2nd day (Fig 1). And then we compared the miRNA expression profiles between the 8th and 5th day. Top 5 changed miRNA in each paired group are listed in table 2 and 3. By comparing the microarray data, we found the amount of decreasing miRNA were much more than that of increasing miRNA. Additionally, the increasing amplitudes of up-regulated miRNA were far less than the decreasing amplitude of down-regulated miRNA. Main changed miRNAs profiles were related with platelet activation, degranulation, platelet-derived growth factor receptor signaling pathway and cell differentiation.

**Fig. 1.**
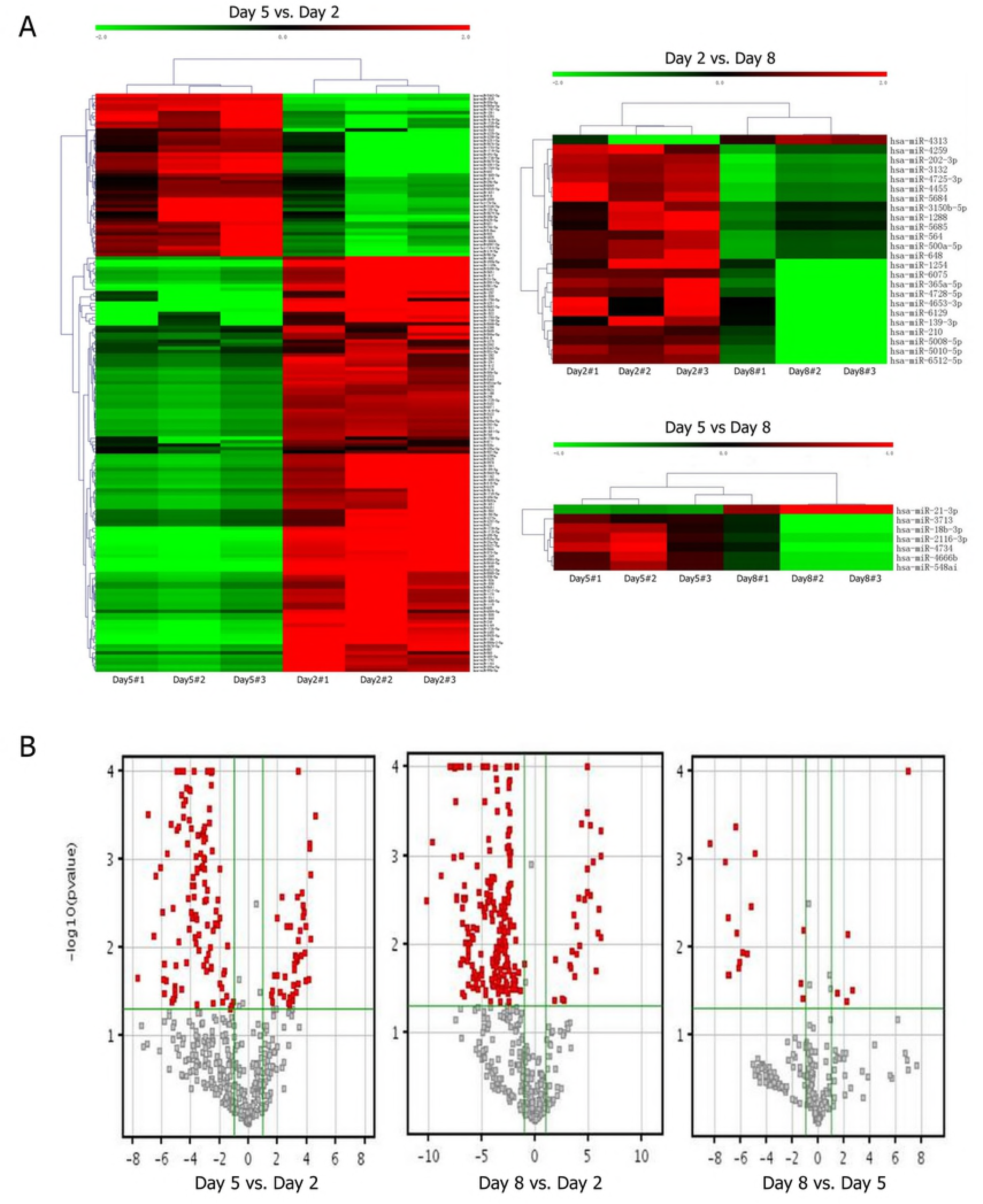
(A) Hierarchical clustering of altered miRNA. Microarray analysis for miRNA expression patterns of platelet Heat-map of deregulated miRNAs that have 2-fold higher (upregulated) or lower (downregulated) Cyanine-3-CTP fluorescence; (B) Volcano ploting microarray analysis showed that have 2-fold higher (up-regulated) or lower (down-regulated) in platelet during storage.

**Table 2.** The miRNAs that exhibited signifcant changes in expression of stored platelet assayed by miRNA Array

| Group | Up-regulation | Down-regulation | Total |
| --- | --- | --- | --- |
| Day 5 vs. Day 2 | 47 | 120 | 167 |
| Day 8 vs. Day 2 | 28 | 202 | 230 |
| Day 8 vs. Day 5 | 5 | 16 | 21 |

Selection criteria of changed miRNAs: Fold change $\geq 2$ , $p < 0.05$ .

**Table 3.**
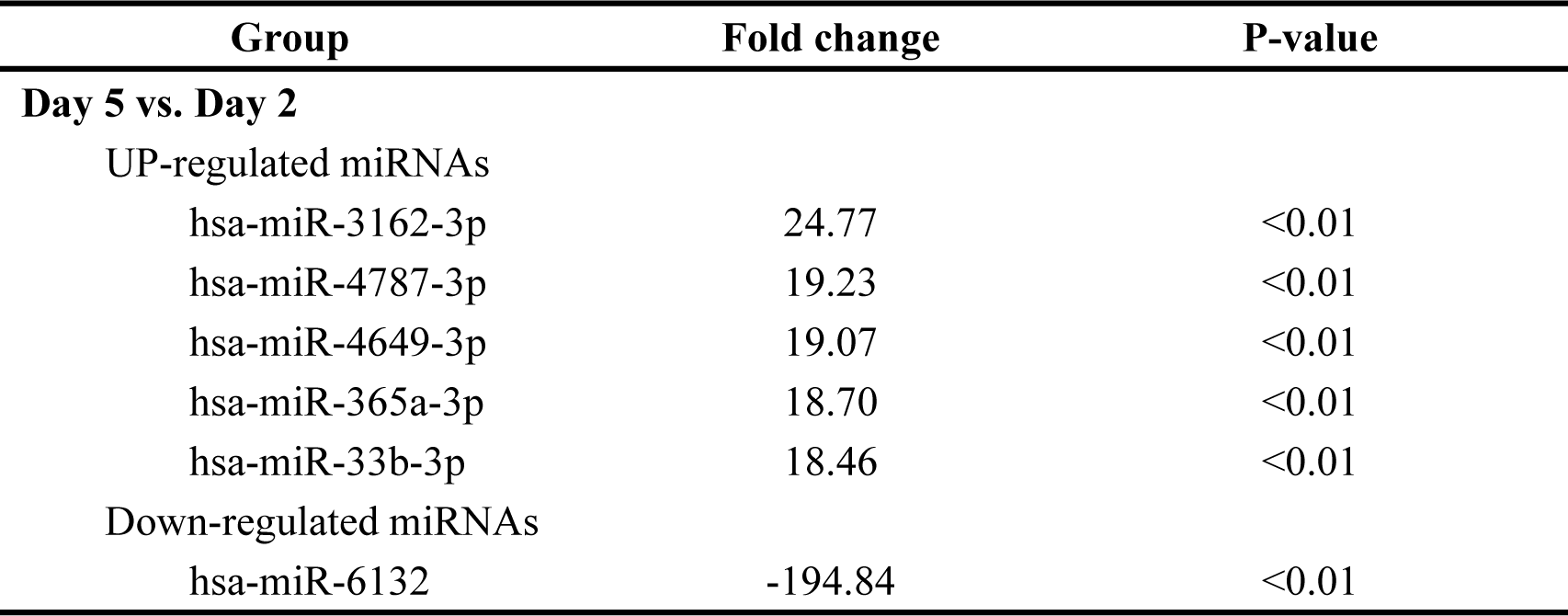
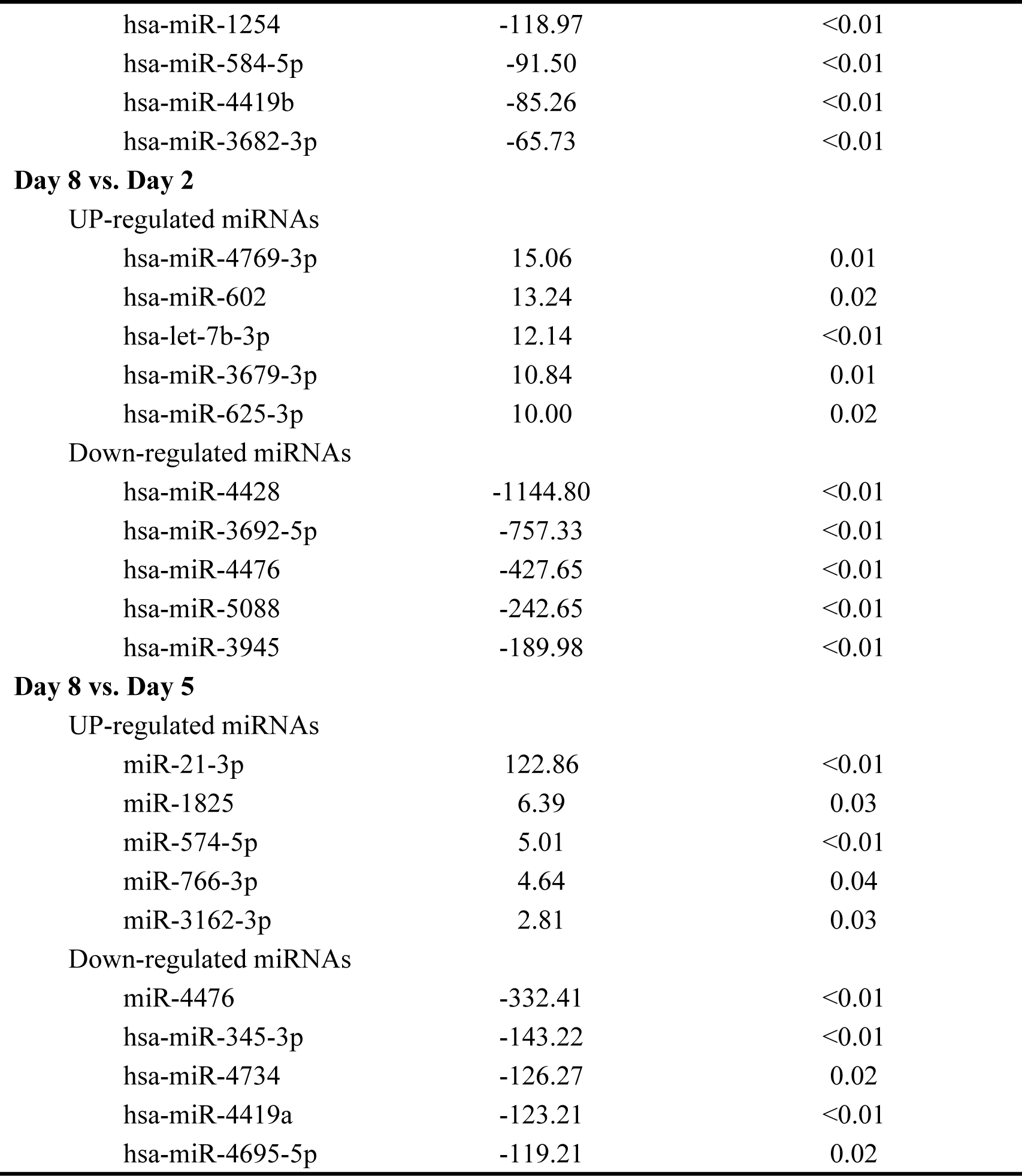
List of Top-5 shifted miRNAs in platelets for different storage duration.

| Group | Fold change | P-value |
| --- | --- | --- |
| **Day 5 vs. Day 2** |  |  |
| UP-regulated miRNAs |  |  |
| hsa-miR-3162-3p | 24.77 | <0.01 |
| hsa-miR-4787-3p | 19.23 | <0.01 |
| hsa-miR-4649-3p | 19.07 | <0.01 |
| hsa-miR-365a-3p | 18.70 | <0.01 |
| hsa-miR-33b-3p | 18.46 | <0.01 |
| Down-regulated miRNAs |  |  |
| hsa-miR-6132 | -194.84 | <0.01 |
| hsa-miR-1254 | -118.97 | <0.01 |
| hsa-miR-584-5p | -91.50 | <0.01 |
| hsa-miR-4419b | -85.26 | <0.01 |
| hsa-miR-3682-3p | -65.73 | <0.01 |
| **Day 8 vs. Day 2** |  |  |
| UP-regulated miRNAs |  |  |
| hsa-miR-4769-3p | 15.06 | 0.01 |
| hsa-miR-602 | 13.24 | 0.02 |
| hsa-let-7b-3p | 12.14 | <0.01 |
| hsa-miR-3679-3p | 10.84 | 0.01 |
| hsa-miR-625-3p | 10.00 | 0.02 |
| Down-regulated miRNAs |  |  |
| hsa-miR-4428 | -1144.80 | <0.01 |
| hsa-miR-3692-5p | -757.33 | <0.01 |
| hsa-miR-4476 | -427.65 | <0.01 |
| hsa-miR-5088 | -242.65 | <0.01 |
| hsa-miR-3945 | -189.98 | <0.01 |
| **Day 8 vs. Day 5** |  |  |
| UP-regulated miRNAs |  |  |
| miR-21-3p | 122.86 | <0.01 |
| miR-1825 | 6.39 | 0.03 |
| miR-574-5p | 5.01 | <0.01 |
| miR-766-3p | 4.64 | 0.04 |
| miR-3162-3p | 2.81 | 0.03 |
| Down-regulated miRNAs |  |  |
| miR-4476 | -332.41 | <0.01 |
| hsa-miR-345-3p | -143.22 | <0.01 |
| hsa-miR-4734 | -126.27 | 0.02 |
| hsa-miR-4419a | -123.21 | <0.01 |
| hsa-miR-4695-5p | -119.21 | 0.02 |

### The confirmation of shifted miRNA using RT-PCR

Our genome wide miRNA screening identified, for first time, many differentially miRNAs expression were regulated during platelet storage. Based on microarray results, miR-21-3p, let-7b-3p and miR-3162-3p, the topist changed miRNAs in each paired group were selected to verify, which were closely related to the function of platelet. The other miRNAs, miR-223, miR-21-5p and miR-155, which have known roles in regulating paletlet functions and large fold changes during storage, were also selected for RT-PCR verification at same time. Real-time PCR assays confirmed the microarray findings (Fig. 2). The expressions of miR-21-3p, miR-21-5p and miR-155 were significantly decreased on the 5th day and 8th day. Meanwhile, miR-223, let-7b-3p and miR-3162 increased significantly on 5th day. These results indicate that, with the increase of bank time, the expression profile of miRNA gradually shift in platelet.

**Fig. 2.**
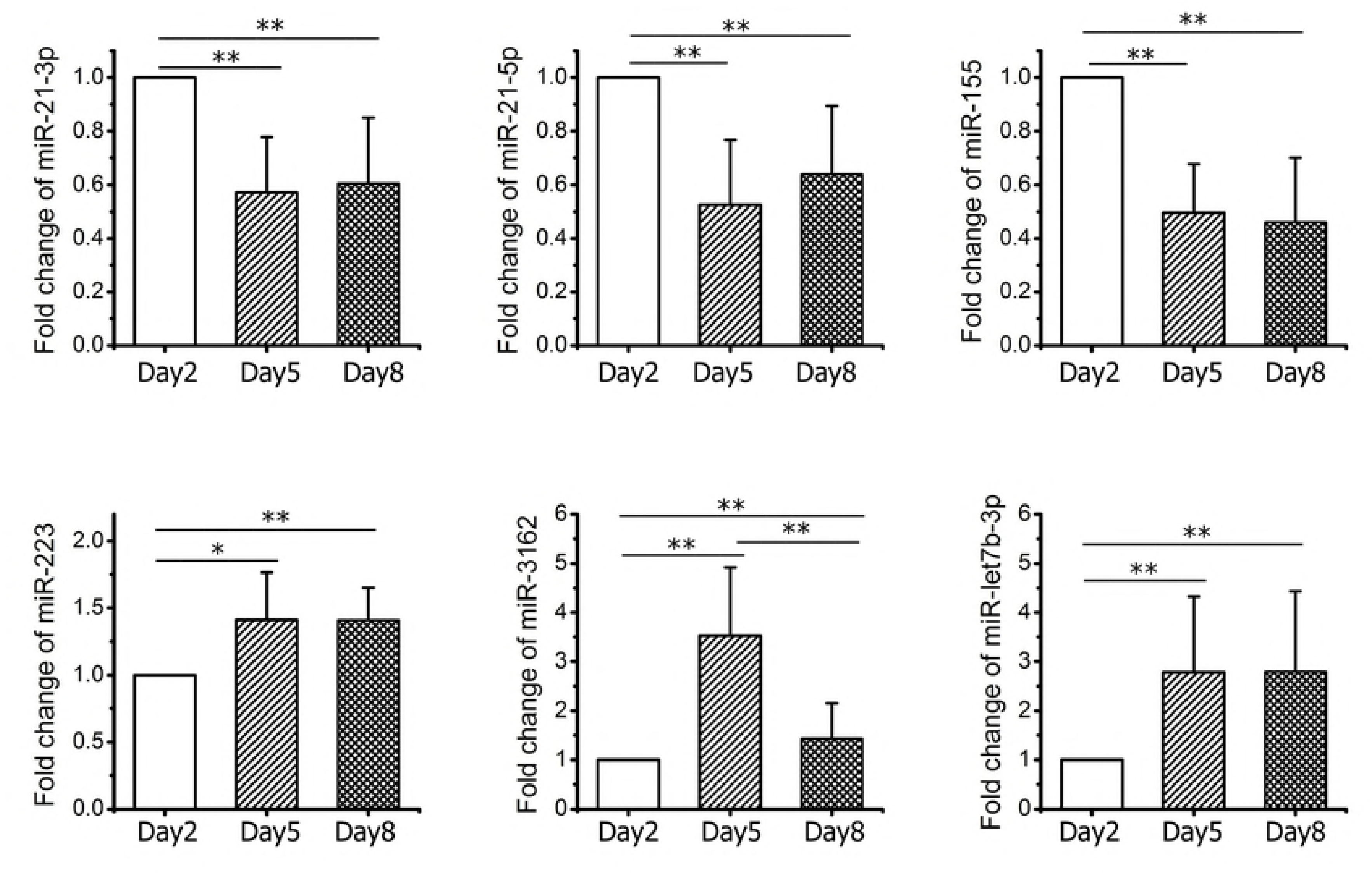
Variation of selected miRNA was tested by RT-PCR in storage platelet. Each bar indicates mean ± SEM (n=15). Data are expressed as the percentage of control and the means ± SD of three separate experiments, each of which was performed in triplicate. \**p* < 0.05, \*\**p* < 0.01.

### Prediction of the signal pathway of platelet activation and aggregation

To assess the biological significance of shifted miRNA level, predictions of the target-related cellular pathways were performed using the KeGG pathways. By KEGG database analysis and related literature reports [22], we predicted the possible schematic representations between miRNA and protein in storage platelet, as shown in Fig 3. These sifted miRNAs were closely related with pathway of self-activation, activation and aggregation. P2Y12, as a specific platelet ADP receptor, played an important role in activation and apoptosis of platelet. The activation of P2Y12 receptor suppressed the Bak/Bax activated by PI3K signaling pathway to inhibit the apoptosis of platelet, and then prevented the abnormity of activating function to maintain normal platelet function [23]. MiR-223 and miR-21 regulated the synthesis of P2Y12 protein by affectting the expression of P2Y12 mRNA. In addition, miRNA-96 and let-7b inhibited their target protein VAMP8 and GPIIIa expression. In this study, miR-223 and let-7b expression levels were found increased during storage, whereas that of miR-21 decreased. The miR-233 and miR-21 might inhibit GPⅡb/Ⅲa and platelet activation process through the regulating expression of P2Y12. But no obvious change was found in miR-96 expression during storage.

**Fig. 3.**
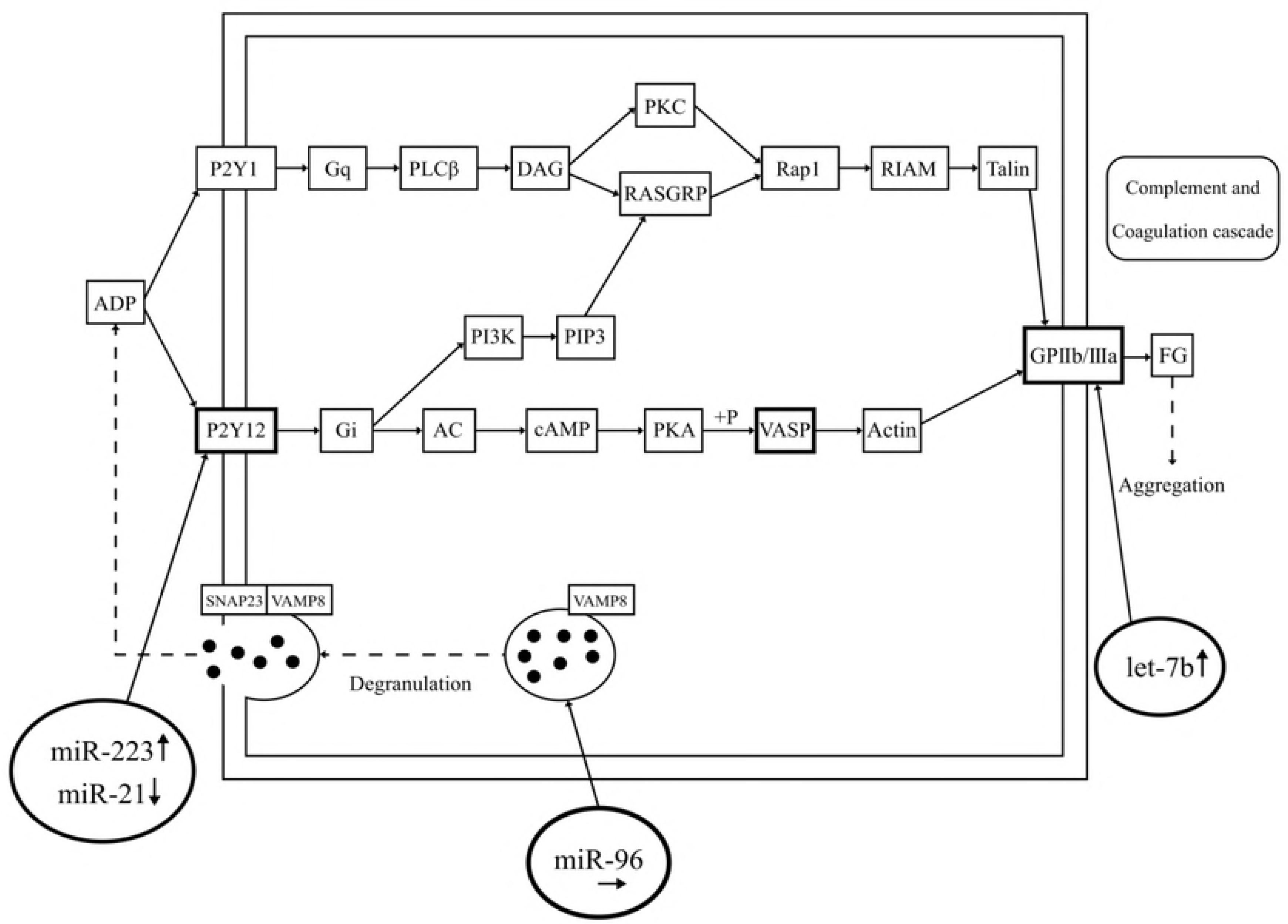
Platelet activation pathway predicted by KEGG database and their possible relationship between miRNAs and platelet agregation.

### The apoptosis of platete during storage

Using flow cytometry we found that with increase of storage time the apoptotic proportions of platete significantly increased. The percentage of AnnexinV-FITC+/PI+ platetes increased in a dose-related manner, reaching a 22.01±1.81% value on the 8th day which was significantly higher than that of platetes on 2nd and 5th day (2.87±0.52% and 22.01±1.81% respectively, *p*<0.01), as shown in Fig 4.

**Fig. 4.**
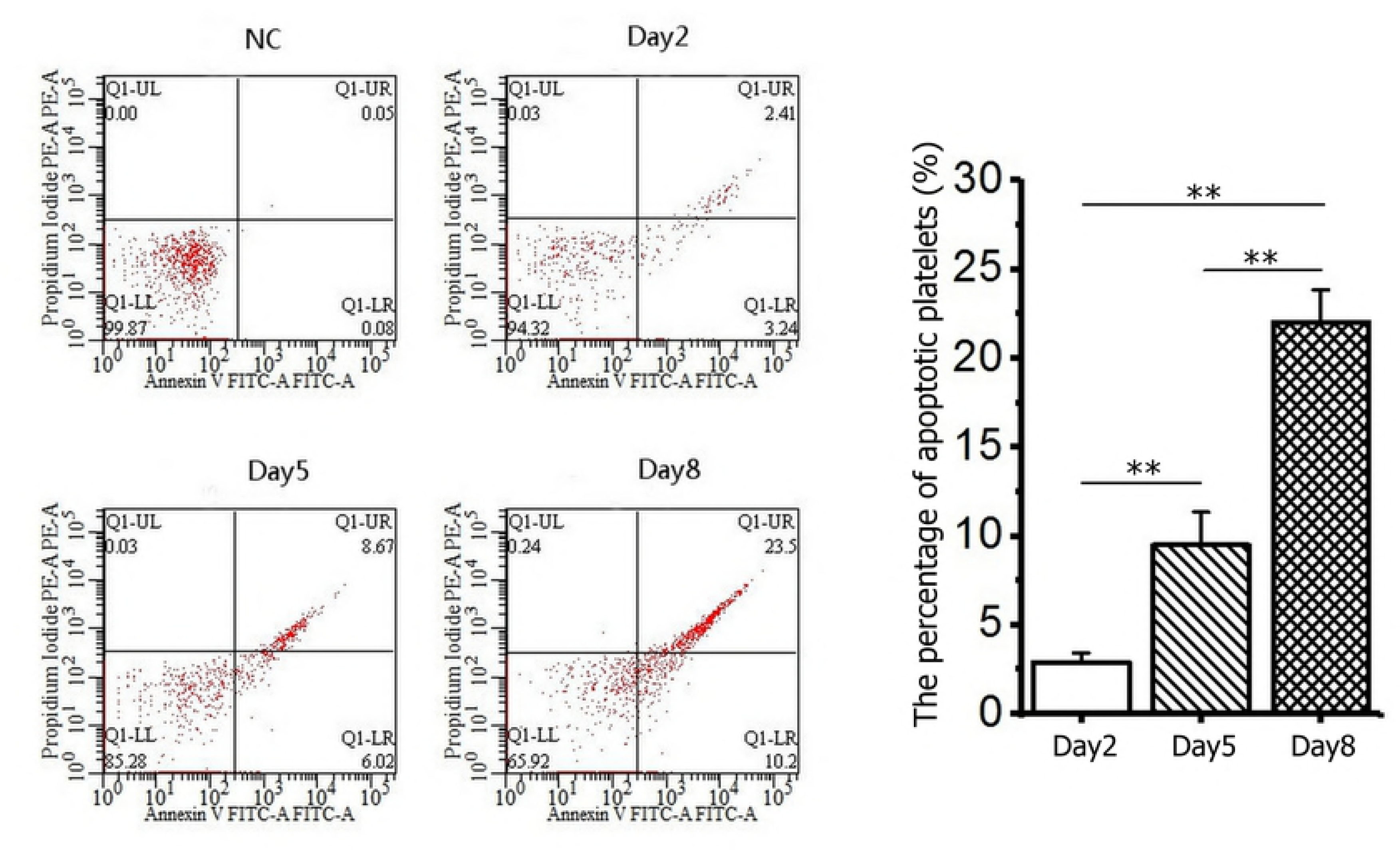
Platelet apoptosis analysis during storage tested by flow cytometry. Data are expressed as the percentage of control and the means ± SD of three separate experiments, each of which was performed in triplicate. \*\**p* < 0.01.

### Platelet reactivity index in storage

Flow cytometric assay was used to determine the VASP phosphorylation level of platelet. Platelet reactivity index (PRI) is closely related with the VASP phosphorylation level, which reflects activation degree of P2Y12. With the increase of storage time, PRI decreased from 72.73% to 18.68% （*p*<0.01）, which indicated phosphorylation levels of VASP decreased, and the activating degree of P2Y12 increased gradually during platelet storage, data were shown in Fig 5.

**Fig. 5.**
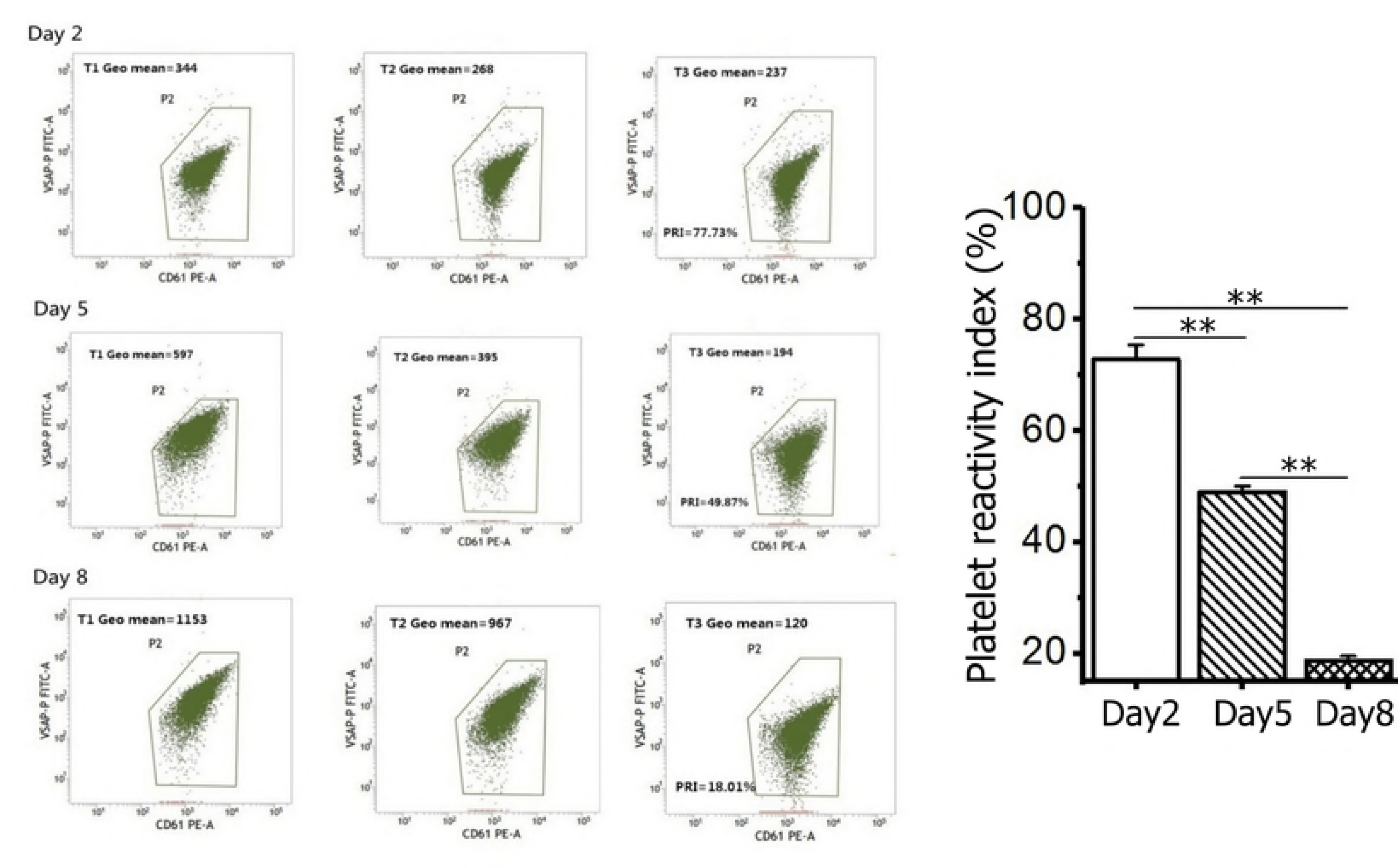
Platelet reactivity index level of platelet concentration tested by flow cytometry. Data are expressed as the percentage of control and the means ± SD of three separate experiments, each of which was performed in triplicate. \*\**p* < 0.01.

### Platelet GPIIb/IIIa expression during storage

By flow cytometry analysis, we found the percentage of CD41a+/CD61+ platelet were 99.16±0.34%, 98.91±0.73% and 98.38±0.74% respectively on 2nd, 5th and 8th day, datas were shown in Fig 6. There was no significantly difference among three groups, which indicates expression of platelet GPIIb/IIIa has not changed, no activation of platelet was detected during the storage.

**Fig. 6.**
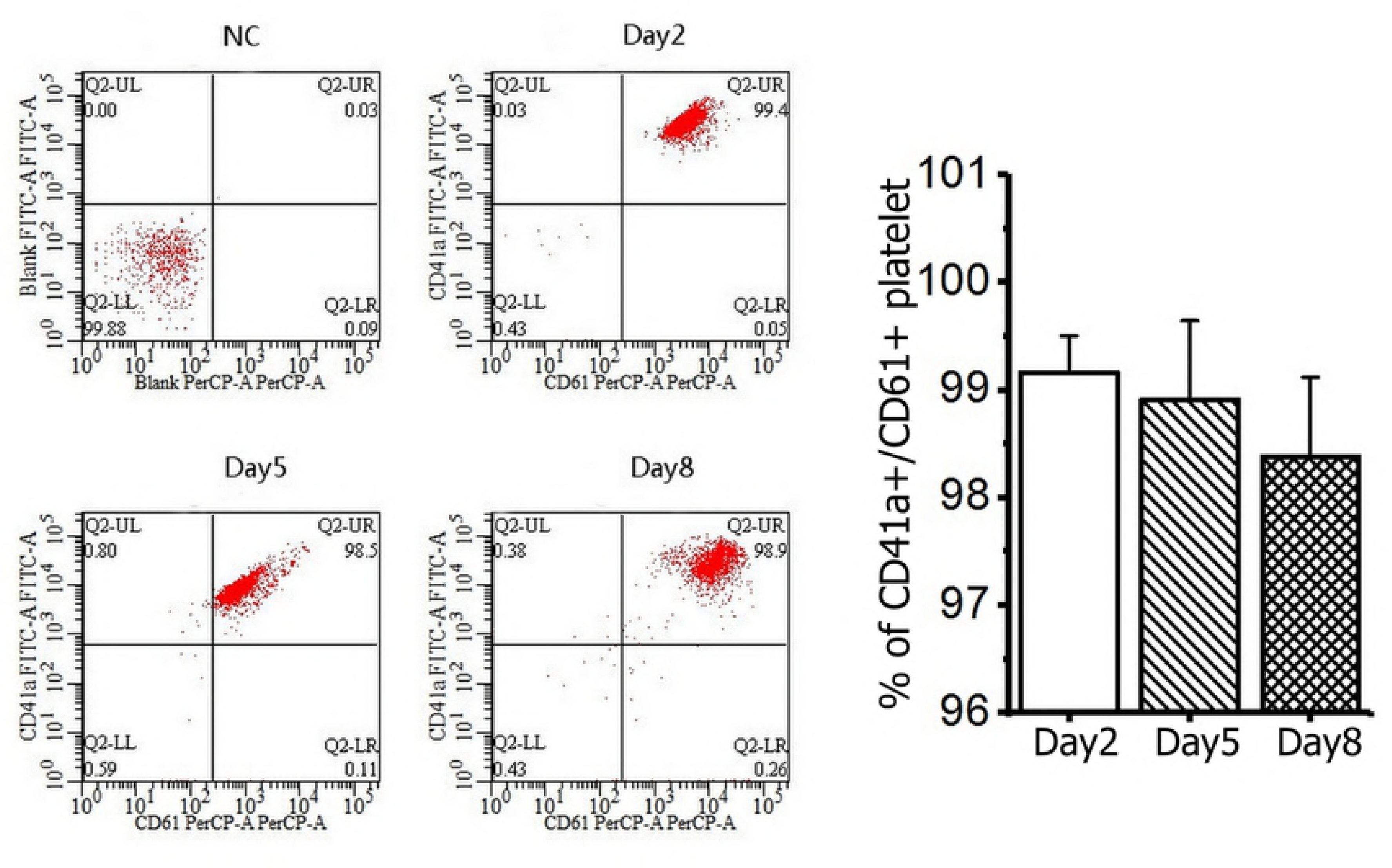
The expression of GPIIb/IIIa in storage platele tested by flow cytometry. Data are expressed as the percentage of control and the means ± SD of three separate experiments, each of which was performed in triplicate.

## Discussion

MiRNAs in human platelets were first reported in 2009 [24]. With the development of detecting and sequencing technology, more and more miRNAs in platelets will be reported. Some miRNAs were found to be closely related to the activity of platelet, mRNA expression protein synthesis and regulation of activation [25, 26]. In this study, by miRNA array detection we identified a large number of miRNAs in stored platelets in vitro. Although the concrete regulating mechanism between miRNA and platelet signaling pathways is not clear, we can speculate the function of miRNA and predict the target-related signaling pathways through bioinformatics database. By KEGG database analysis and related literature reports [22], the possible schematic representation between miRNA and protein in platelet was predicted.

We found miR-223 regulated the expression of P2Y12 mRNA by affectting the synthesis of P2Y12 protein and this process was also adjusted by miR-21. Our results show that expression of miR-223 increased during storage, while the expression of miR-21 reduced. By flow cytometry we further confirmed the effect of altered miRNA on related proteins and the activation of P2Y12. P2Y12, an important part of the activation and aggregation of platelet, could activate final target protein GPⅡb/Ⅲa by regulating the aggregating signaling pathway [23, 27]. Therefore, we speculated that miR-223 and miR-21 affected the functions of GPⅡb/Ⅲa by P2Y12 protein. We found GPⅡb/Ⅲa, the marker of platelet activation, show no obvious change during storage, this indicated the activation and aggregation of platelet is not significant. Furthermore, the unaltered GPⅡb/Ⅲa level might be related to bidirectional regulation of miRNA to GPⅡb/Ⅲa. For example, the increasing miRNA let-7b inhibited targeted gene of GP Ⅲa to block the platelet activation and maintain their normal function. The miR-233 and miR-21 inhibited GPⅡb/Ⅲa and platelet activation process through the regulation of P2Y12.

In this study, we found there are many alterd miRNAs, including miR-21-3p, miR-21-5p, miR-155, miR-223, miR-3162-3p, let-7b-3p, but most of these expression alteration remains not to be elucidated. MiR-21 is the key factor regulating proliferation and migration of human vascular smooth muscle cell and can be induced by platelet-derived growth factor, which may become an attractive treatment method of proliferative vascular diseases [28]. Our results show that the expression of miR-21-3p and miR-21-5p had decreased significantly, especially on the 5th day on which it declined over 40%. This indicated that the infusion of platete stored for 5 days could lead to decreasing expression of nuclear transcription factor activator protein-1 and alpha smooth muscle actin. MiR-155 is reported to contribute to proliferation of the megakaryocyte and inhibit its differentiation [29]. The decline of miR-155 expression in platelet storage perhaps implied the continuation of the decline of megakaryocyte, which is likely to maintain the platelet function as a negative regulator. In this study, let-7b expression levels were found increased during storage, which could inhibit their target protein expression of VAMP8 and GPIIIa [24].

The predicted data show that P2Y12 plays an important role in aggregation and activation of platete and is closely related to the miR-223 expression. The P2Y12 receptor was first described in blood platelets as Gi-coupled ADP receptor, and it is the key to the complex processes of activation and aggregation [23]. During storage, miR-223 and P2Y12 expression level changed significantly and they were mutually controled, which is also associated with the activation of platelet aggregation. Meanwhile, the expression of miR-223 in apheresis platelet was very high and easy to be detected, which is more likely to be a biological marker of platelet storage. The normal life of platelet is 8~11 days [3]. Rinder confirmed that platelet can still live for 5-7 days in vivo after being stored in vitro for 5 days [30]. Although the preservation duration of platelets in vitro is still the 5 day of according to international standard, with the improvement of apheresis platelet storage technology and storage conditions, shelf life of platelets is expected to be prolonged in vitro. MiR-3162 is thought to be involved in regulating phenotype of regulatory T cells in rheumatoid arthritis patients, and its subunit miR-3162-3p was first found to be associated with platelet function. MiR-96 is reported to inhibit the expression of VAMP8, which plays an important role in platelet activation pathways by ADP [31], but we did not found any changes of miR-96 expression in storage platelet. By the bioinformatic analyses and the GO analysis tool, we found there are total 167 altered miRNA on the 5th day, including 115 target genes associated with platelet activation, 40 target genes closely related with platelet degranulation, 15 target genes associated with platelet-derived growth factor receptor signaling pathways, and 7 target genes associated with stem cells. This indicates that there are many mysteries to be discovered about miRNA and their rgulations.

In summary, this study showed many miRNAs were found altered expression levels in banked platelet. Variational miRNAs expressions levels are related to proteins about platelet aggregation, such as P2Y12，VASP，GPⅡb/Ⅲa. Through the KEGG database and the RT-PCR validation, miR-223, miR-21 and let-7b can influence the pathway of platelet aggregation. The miR-223 is easy to detect and showed altered expression level increased sharply during storage, and miR-223 has the potential to be the biomarker of storage lesion. More researches on miRNA and their target genes interaction and regulation network mechanism need to be elucidated.

## Acknowledgments

The Project Sponsored National Natural Science Foundation of China (81702058).

## Authorship contributions

Zhaohu Yuan, Qianyu Wu and Xiaojie Chen, performed the research, analyzed data, and wrote the report. Yaming Wei designed the research study and analyzed the data.

